# Pollen meta-barcoding reveals foraging preferences of honeybees (*Apis mellifera* L.) along space-time gradient in Japan

**DOI:** 10.1101/2021.08.05.455320

**Authors:** Grégoire Noël, Arnaud Mestrez, Philippe Lejeune, Frédéric Francis, Masayuki Miwa, Koichi Uehara, Ayako Nagase

**Author notes:** These authors contributed equally to this work (Grégoire Noël, Arnaud Mestrez).

## Abstract

The availability of pollen in urban-rural landscapes is an essential factor that influences the population dynamics of insect pollinators. The amount and diversity of pollen play a pivotal role in the foraging ecology of pollinators for their growth and health, but investigations on the spatio-temporal patterns of foraged plants remain rare, especially in cities as neo-ecosystems. Here, we explored the temporal foraging habits of a highly polylectic pollinator (*Apis mellifera* L.) in a study area, including different landscape classes from rural to urban areas. Mixed-pollen in each month and each location (N = 17) were analysed using DNA meta-barcoding to identify plants visited by honeybees. The results showed that the landscape class (rural, suburban and urban areas) explains spatial variations in the plant composition foraged by honeybees, but not in taxa richness. Furthermore, pollen diversity and plant composition showed a strong seasonal dependence. Furthermore, a higher plant richness and foraged woody taxa was found to occur in spring, which was mainly dominated by the genera *Prunus* and *Acer*. In summer and autumn, the genera *Trifolium* and *Plantago* of the herbaceous stratum were the most visited plants. The Fabaceae, Rosaceae, Brassicaceae, Plantaginaceae, and Onagraceae plant families were the most frequently observed in all combined samples. The present study contributes to a broader understanding of the ecology and floral preferences of honeybees, on which urban planning can rely to promote biodiversity in cities.

## Introduction

Ongoing urbanisation is one of the main drivers of landscape degradation and pollinator biodiversity loss [1–5]. Indeed, floral resources are becoming scarcer under the pressure of urban fragmentation, and the increase in impervious surfaces is rendering nesting sites inaccessible to pollinators [6,7]. However, recent studies have revealed that cities can also act as a refuge for pollinators [8], particularly for bees [9]: (i) cities are less exposed to pesticides [5,10], (ii) urban management sustainably permits the maintenance of their floral resources [11,12], and (iii) urban areas are configured with a heterogeneity of green spaces, which would be favourable to the foraging preferences of bees [13]. Moreover, flowerbeds in the urban matrix are highly attractive and are a source of pollen and nectar for insect pollinators [14,15]. Therefore, plant selection in greening projects must be pollinator-friendly. With an increasing popularity in beekeeping activities in cities, honeybees (*Apis mellifera* L.) contribute to urban plant pollination, generate profits of by-products, and provide environmental education [16,17]. However, the massive introduction of urban honeybees has led to growing concerns about detrimental effects on wild pollinators through an increase in floral resource competition and the spillover of shared pathogen agents [18,19].

As a eusocial species, honeybees organise their floral resource collections (i.e. nectar and pollen) through a complex communication system within their colonies. According to plant phenology, honeybee scouts rapidly recruit their siblings to forage rich new patches of flowers using a characteristic waggle dance [20]. The foraging decision-making system of the colony can vary from day to day or within the same day following real-time nectar and pollen availability in the surroundings. Throughout its active seasons, the colony constantly maintains a balance between its biomass and energy management, combined with a diversity of floral sources, to fully benefit from the abundance period, and makes reserves for incoming winters [21,22]. Pollen diversity provides substantial resources in terms of protein, lipid, vitamin, and mineral supplies [23]. Large amounts of pollen (15–30 kg) are collected annually, mainly for brood production during summer [21,24,25]. The quality and diversity of pollen are also essential for strong health and better immunity, as well as longevity and parasite tolerance of the bees and the colony [26–29].

The preservation of ecosystem functioning relies on the mutualistic networks of pollinators and plants. Several methods are used to assess these interactions: the observation of floral visits, digital tracking systems to capture floral visits, chemical signatures of pollen, pollen genetic sequencing, and pollen light microscopy [30]. Identifying a pollen species or genus by light microscopy from mixed pollen samples, also known as melissopalynology [31], is a time-consuming process, even for well-trained experts [32] that results in low taxonomic resolution, usually at the family or genus rank at best [33–36]. With the advent of high-throughput sequencing (HTS) techniques, DNA meta-barcoding approaches have become reliable methods to obtain faster taxonomic profiles with higher resolution of mixed pollen collected from bees or flowers [37–39]. To elucidate the plant taxonomic composition of mixed-pollen samples, the meta-barcoding process can be based on different genetic markers, such as the *rbcL*a, *matK, trnH-psbA, trnL*, and *ITS* regions (mainly *ITS2*), which require high inter-specific and low intraspecific variability [36,39–42]. These selected loci and the primer set used for amplification drive the range of taxonomic inferences [33]. The identification and quantification of plants visited by pollinators is an ecological issue for natural resource management and preservation in urban environments [36,43]. Indeed, the fragmentation of urban matrix usually leads to the creation of small, remote, and intensely maintained green spaces [44], which could influence the foraging distance of honeybee workers in terms of pollen richness and quantity [45]. The composition of agricultural or urban landscapes heavily impacts the foraging preferences of honeybee workers [46,47]. As the most polylectic bee forager [48], honeybees can mitigate the degradation of floral resources by enlarging their foraging area [49,50]. Moreover, seasonal shifts greatly impact the pollen availability for honeybee colonies, according to the phenology of the floral resources [51,52]. In temperate climates, the foraged plant traits also vary according to the course of the seasons: spring is dominated by trees and shrubs, summer has more herbaceous species, and autumn is characterized by woody vines [52]. Nonetheless, the space-time effects combined with plant traits in the foraged plant community have yet to be studied extensively [53].

The present study aimed to determine the floral preferences of *A. mellifera* to promote pollinator-friendly plant biodiversity through integrated urban greening projects. To achieve this aim, the taxonomy of pollen foraged by honeybees was identified over the seasons along an urban-rural gradient from different locations in the Kanto region, Tokyo, and its surroundings in Japan. The research questions addressed were as follows: (i) How does the composition of the foraged flower community (and the foraged plant traits) vary according to an urban-rural gradient? (ii) How does the composition of foraged flower communities (and foraged plant traits) vary according to the course of the seasons? Lastly, the pollen availability of plant taxa composition foraged by honeybees was evaluated.

## Material and methods

### Study area and experimental set-up

We selected 17 apiary locations, with homogeneous climatic and altitudinal conditions (Table S1), distributed along the urbanisation gradient in the Kanto region of Japan (Fig 1). One hive per apiary was used for pollen sampling. From March to September 2019, pollen samples were collected using pollen traps at the entrance of the same hive (Fig S1). This experimental setup was used to collect pollen balls from the hind legs of honeybee foragers with a standardised honeybee-size mesh and tray [54]. Then, the contents of the pollen traps were discharged into labelled 50-ml conical tubes and stored at −20°C. However, due to meteorological conditions and the personal schedules of the beekeepers, the sampling date, collection frequency, and operational time of the pollen trap varied from site to site. For subsequent analysis, the sampling dates were discretely pooled by month and analysed in R version 4.0.2 [55]. We used Spearman’s rank-order correlation between the number of taxa per sample and the sampling length in hours to test whether the data could be treated independently of the sampling length. All samples were sent to the private company, Bioengineering Lab. Co., Ltd. (https://www.gikenbio.com/, consulted on 20/07/2020) for the processing of the meta-barcoding of the mixed pollen samples.

**Fig 1.**
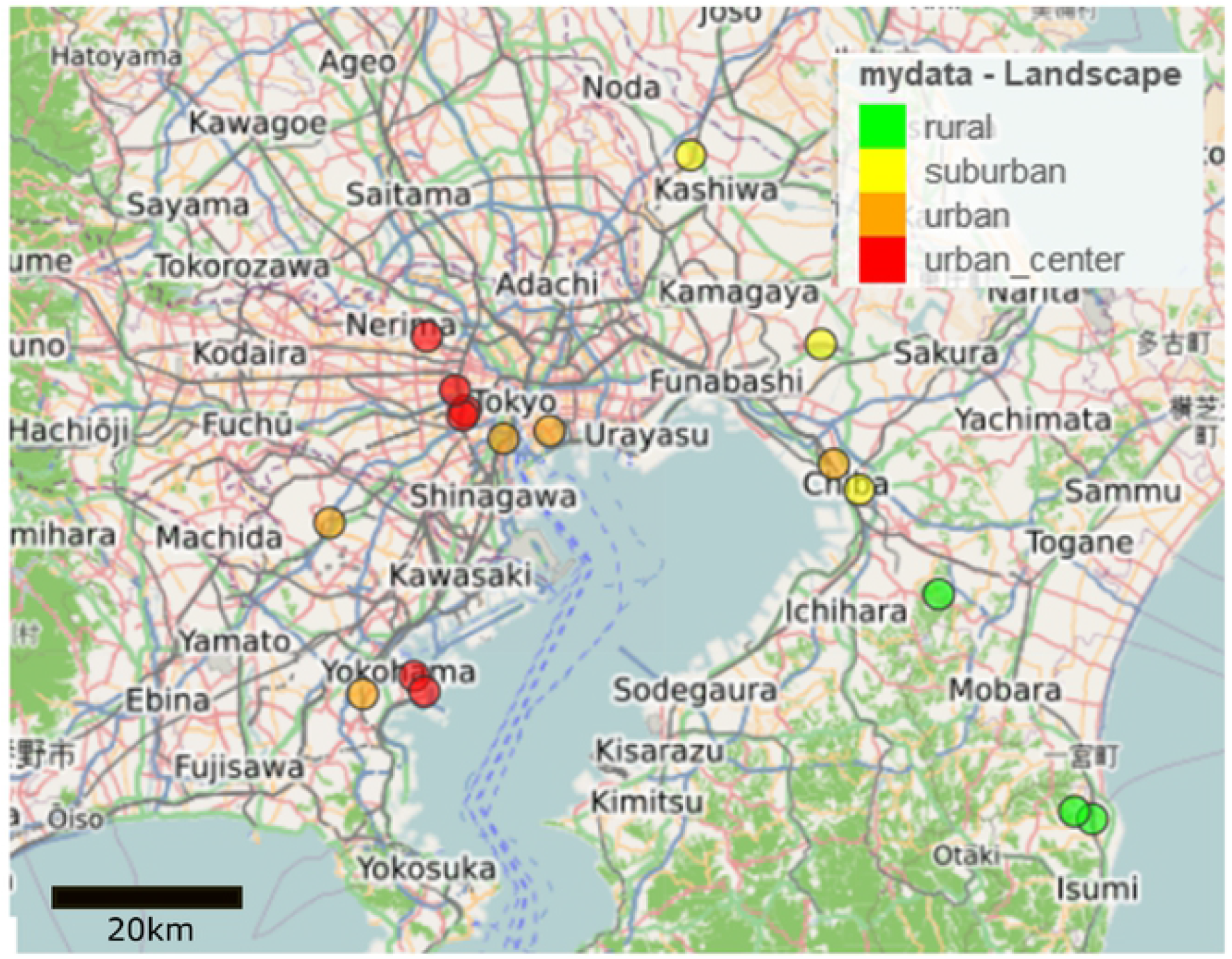
Selected hive locations along Tokyo bay (Japan). Each colour corresponds to the landscape type resulting from the cluster analysis of the study sites based on k-means approach. The map was drawn using *Openstreetmap France* from *mapview* in R [56].

### Landscape analysis

Using remote sensing techniques, the landscape structure was investigated within a 6-km radius around each hive location (Table 1); this distance enclosed 95% of the forage area per colony [21]. With the help of Planet Labs Inc. (2020) [57], we used multi-spectral images (RGB, NIR) with 3-m pixel resolution. To fully exploit the potential of the data, the cloud cover condition was set to a maximum of 5%. Planet data are relevant for computing and mapping high-resolution terrestrial above-ground vegetation at the landscape scale [58]. For each planet image, the normalised difference vegetation index (NDVI) was computed using the red and near-infrared bands based on band rationing, which allowed for the delineation of the vegetation cover from other types of land cover [59]. Classes were created with the function *reclassified* from the *raster* package in R [60] by defining the NDVI threshold value to 0.2 to distinguish the non-vegetation (NDVI: from −1 to 0.199) from the vegetation (NDVI: from 0.2 to 1) [61,62]. A majority filter, with a 6 × 6 filter kernel size, from the *whitebox* package in R [63], was applied to smooth the result and aggregate regions of high uncertainty. Landscape classifications at the site level were performed using demographic data (i.e. number of inhabitants per admin units) and landscape metrics from the *lconnect* [64] and *landscape metrics* [65] packages in R. To classify our sites along an urban-rural gradient [44,66], we retained selected data: number of inhabitants per km^2^ (dpop), the integral index of connectivity (IIC) [67], the effective mesh size (MESH) [68], Shannon’s evenness index (SHEI) [69], the vegetation cover proportion (veg cover), vegetation patch density (Pd) [70], and the median vegetation class NDVI (NDVI median) (Table 1).

**Table 1.**
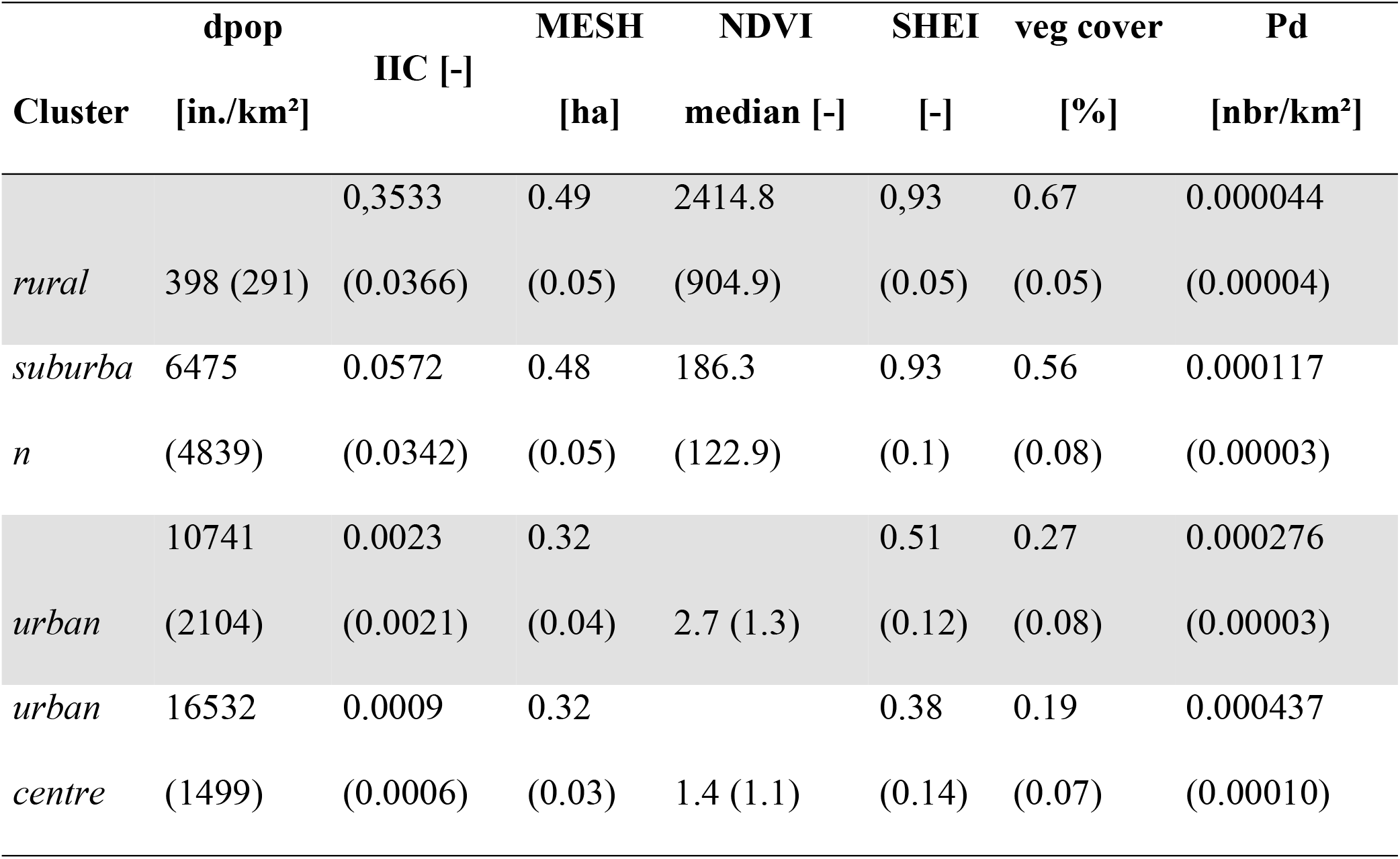
Mean and standard deviation of variables among the landscape classes. The units of landscape variables are given in square brackets. The standard deviation is provided after the mean.

We conducted principal component analysis (PCA) of the landscape dataset to visualise the differences among our study sites. The unsupervised k-means clustering method was applied to delineate the landscape category along the urban-rural gradient into k groups. Before initiating the analysis, the data were standardised using the *scale* function in R to make variables comparable. As a result, the clustering algorithm was independent of any variable unit. The number of k groups required to be defined as the first step was determined using the elbow method [71]. The k-means partitioning analysis was performed using the *k-means* function with 25 random sets [72] and the *factoextra* package in R for PCA graphical representations [73].

### Molecular techniques

#### DNA extraction

First, pollen samples (0.5 g) were lyophilized using a lyophilizer freeze dryer VD-250R (TAITEC, Koshigaya, Saitama, Japan). After being ground at 1500 rpm for 2 min using a ShakeMaster NEO homogeniser (bms, Shinjyuku, Tokyo, Japan), DNA was extracted using the protocol of MPure Bacterial DNA Extraction Kit (MP Biomedicals, Irvine, CA, USA). DNA purification of the samples was performed using the MPure-12 Automated Nucleic Acid Purification System (MP Biomedicals, Irvine, CA, USA). Quality control of DNA extracts was conducted using Synergy H1 (BioTek, Winooski, VT, USA) and QuantiFluor dsDNA System (Promega, Madison, WI, USA).

#### Library preparation and DNA sequencing

Libraries were produced using a 2-step tailed polymerase chain reaction (PCR) method. The first PCR amplification was conducted using internal transcribed spacer (ITS1) primers designed by Masamura [74], coupled with MiSeq-specific adapters and Illumina index sequences. The second PCR amplification was conducted using index primers. PCR reactions were carried out in a reaction volume of 10 μL containing 1.0 μL of 10× Ex Buffer, 0.8 μL of nucleoside triphosphate dNTPs (each at 2.5 mM), 0.5 μL for both forward and reverse primer at a concentration of 10 μM, 2.0 μL of DNA template normalized at 0.5 ng/µL, 0.1 μL of DNA polymerase ExTaq at 5 U/μL (TaKaRa, Otsu, Shiga, Japan) and 5.1 μL of double-distilled water. The PCR profile was as follows: 2 min of denaturation at 95°C, followed by 10 cycles with 30 s of denaturation at 95°C, 30 s of annealing at 57°C, 30 s of elongation at 72°C, and a final elongation at 72°C for 5 min. The PCR products were purified using AMPure XP (Beckman Coulter, Brea, CA, USA). Library concentrations were determined using a Synergy H1 microplate reader (BioTek, Winooski, VT, USA) and a QuantiFluor dsDNA System (Promega). Library quality was evaluated using a fragment analyser (Advanced Analytical Technologies, Ankeny, IA, USA) with a dsDNA 915 Reagent Kit (Agilent, Santa Clara, CA, USA). The generated library was sequenced using MiSeq Illumina technology (Illumina, San Diego, CA, USA) through a 2× 300 paired-end run.

#### Data processing

We used “FASTX Barcode Splitter” from Fastx toolkit, a short-reads pre-processing tool, to extract only the Miseq reads with sequences readings that matched the primers used exactly [75]. Next, the reads were denoised and filtered using Sickle software [76] with an overlap quality value of 20. Trimmed reads with fewer than 150 bases were discarded. The remaining reads were merged using FLASH (version 1.2.11) paired-end merge script [77] using the following conditions: fragment length after merging of 420 bases, read fragment length of 280 bases, and minimum overlap length of 10 bases. The paired readings were aligned with the reference sequences using the open-source bioinformatics pipeline Qiime 2.0 [78]. A Qiime workflow script was used for OTU creation and taxonomic assignment. A naïve Bayesian classifier was used for taxonomic assignment with the output sequence [79,80]. Following the taxonomy classification step, taxa-abundance data and operational taxonomic unit (OTU) data were applied to the R environment [55]. First, the assignment of all OTUs below the identity threshold of 97% was discarded [46,81]. Next, the number of reads was sorted by genus and sample (i.e. site and collection date), and was then expressed as the ratio between the read count and the sum of read number per sample for each genus. Genera accounting for less than 0.05% of the total number of readings for a single sample were excluded to prevent false positives. Moreover, a sample was removed if it accounted for less than 1000 reads to limit inferences from insufficient sequencing depth [82].

### Taxonomic analysis

Given the poor correlation between the actual pollen proportion per foraged plant taxa and the sequencing read proportions from one locus [36,41], the analyses undertaken were based exclusively on the incidence (i.e. presence/absence binary arrays) dataset. This process prevents the misinterpretation of DNA metabarcoding data [83]. The richness of the pollen samples (i.e. the number of distinct taxa of foraged plants) was analysed as a function of the months and landscape classes. Differences were assessed using analysis of variance (ANOVA) on richness (response), across independent landscape classes (between-subject factors), and along time (within-subjects factor). Prior to the test, the data were mathematically root-squared transformed to respect the normality (Shapiro–Wilk test) assumption for each combination of factor levels. Then, the assumptions of homogeneity of variance (Levene’s test), homogeneity of covariances (Box’s M-test), and sphericity (Mauchly’s test) of the landscape classes were verified at each level of the time variable. Finally, Tukey’s post-hoc test was applied to analyse the pairwise differences. For multivariate analysis, the pollinated plant taxonomic composition of the samples was studied across sites, sampling periods, and landscape classes using the Jaccard dissimilarity metric from the *vegan* package in R [84]. This asymmetric distance coefficient addresses the problem of double zero, which is essential when studying data on community composition along a gradient. Differences in plant composition between sampling periods and landscape classes were investigated by permutation-based multivariate analysis (n = 999) of variance using the *adonis* function [85]. If the PERMANOVA results were significant, a post-hoc multilevel pairwise analysis with Bonferroni correction was performed using the *pairwiseAdonis* package in R [86]. The dissimilarities in the structures of pollinated plant communities were displayed using non-metric multidimensional scaling (NMDS) with 999 permutations. All graphics were generated using the *ggplot2* package in R [87].

### Indicator species and trait-based analysis

Similarity percentage (SIMPER) analysis was applied to identify how the taxonomic composition differed from the environmental conditions (landscape type) and changes (season). This step allowed us to identify the sampled taxa that contributed significantly to the dissimilarities among the months or landscapes. Finally, to analyse the foraging preferences of honeybees concerning plant traits, each identified taxa in the samples was characterised by its nature, including herbaceous (no woody stems above ground) or woody taxon (tree, shrub, liana), and its native status, including native, alien, or cultivar taxa. The plant trait database was built using information from Ylist [88] and ©Species2000 [89] for the Japanese plant dataset. To determine if the proportion between the different traits varied with the seasons and landscape types, the G.-test of independence for contingency table was performed using the *RVAideMemoire* package in R [90]. The G-test is based on the log likelihood ratio and tests whether the relative proportions of one categorical variable (i.e. plant nature or nativity status) are independent of the second categorical variable (i.e. season or landscape). Next, post-hoc pairwise comparisons were conducted between pairs of proportions using the Bonferroni correction of the p-values [91].

## Results

### Landscape classification

The method of differentiating vegetation from impervious surfaces using NDVI provided convincing results after crosschecking, even in the complex environment of an urban matrix. The two first dimensions of the landscape PCA from landscape variables of our sampling sites described a high percentage of the variance (axis 1 = 81.6% and axis 2 = 11.0%; Fig 2). According to the elbow method (Fig S2) of the k-means partitioning, we classified our sampling sites into four landscape classes according to a rural-urban gradient: rural (n = 3), suburban (n = 3), urban (n = 5), and urban centre (n = 6). The urbanised locations were driven by a much higher demographic density compared to the other landscapes (Fig S3). Moreover, following the decrease in the proportion of vegetation along the rural-urban gradient, it can be assumed that the higher patch density in the cities was induced by the presence of many smaller plots, such as private garden patches. In contrast, the rural sites demonstrated a higher connectivity between the patches. Finally, suburban landscapes occupied an intermediary position between rural and metropolitan areas.

**Fig 2.**
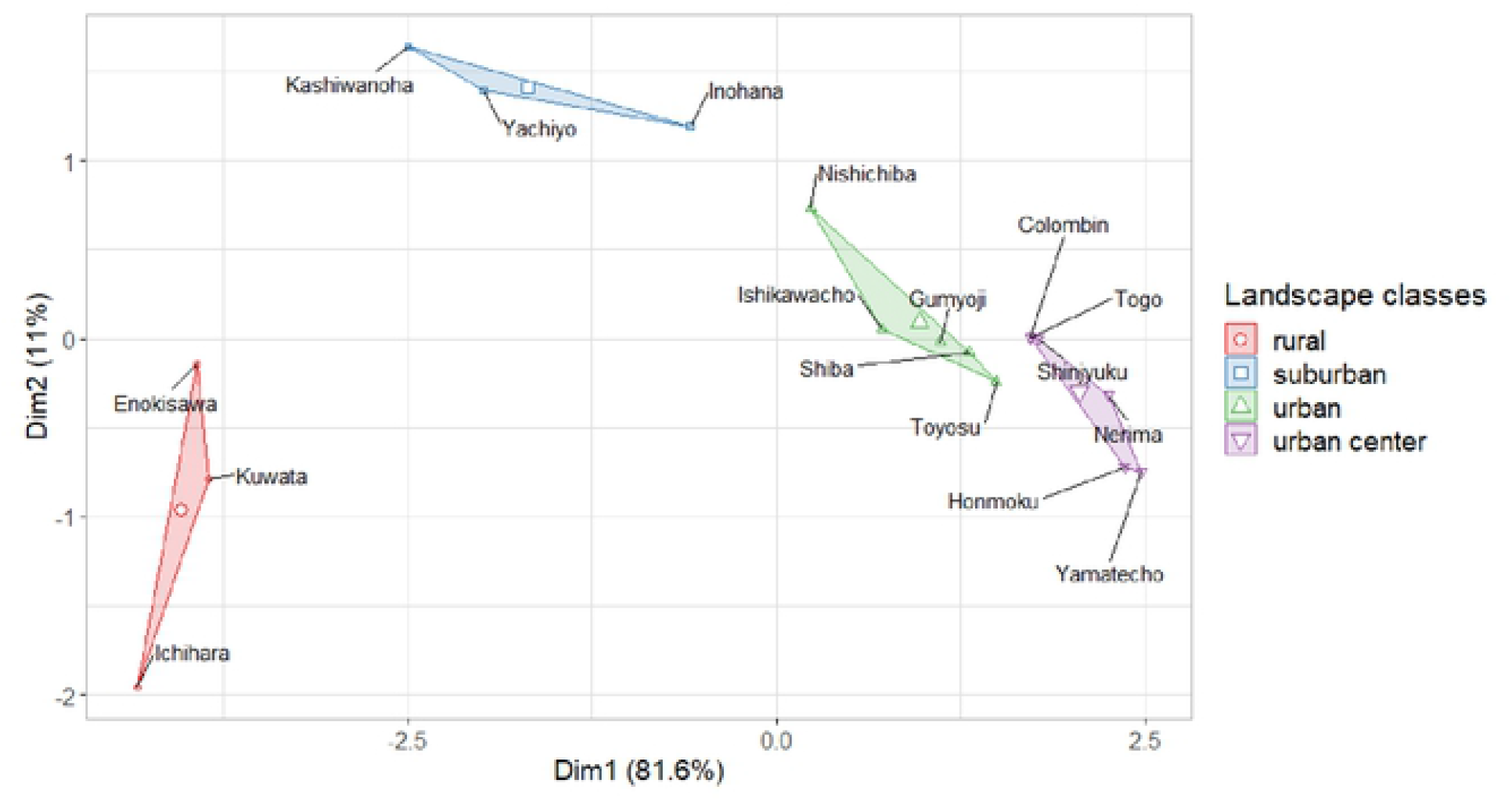
Landscape cluster analysis of selected locations based on k-means approach. The axes represent the first two principal components of the PCA analysis. The dot shapes and colours represent the resulted landscape classes: circle/red = rural landscape; square/blue = suburban landscape; triangle/green = urban landscape; reversed triangle/purple = urban centre landscape.

### Taxonomic analysis

Illumina sequencing generated a total of 8,179,602 paired-end raw reads for the 143 pollen samples from the 17 sites throughout the 7 months of pollen sampling. After assembling and filtering, 6,799,314 reads (83.2%) were obtained for analysis, with a mean count of 47,548 ± 27,464 (SD) reads per sample. By taxonomic assignment of the meta-barcoding dataset, we identified 307 plant flower taxa from 74 families and 187 genera. Prior to the analysis, the richness was not correlated with the sampling length, showing a very weak relationship (*rs* [143] = −0.17, *p <* 0.05), allowing us to consider the statistical independence of all our pollen samples.

Plant richness ranged between 3 and 42 pollinated plant taxa per sample, with an average of 12 (*SD* = 6.2). Mixed ANOVA was not able to detect a significant interaction between landscape type and collection time on the squared root taxa richness (*F* = 0.95; *p* = 0.52). The follow-up tests determined that the landscape type implied no statistically significant differences (*F* = 2.00; *p* = 0.12), but the pollen richness (squared root plant taxa richness) varied significantly across the months (*F* = 5.22; *p <* 0.001). The early spring (March and April) showed significantly higher pollen diversity (Tukey’s test: *p <* 0.05) than the late growth season (July, August, and September) (Fig 3).

**Fig 3.**
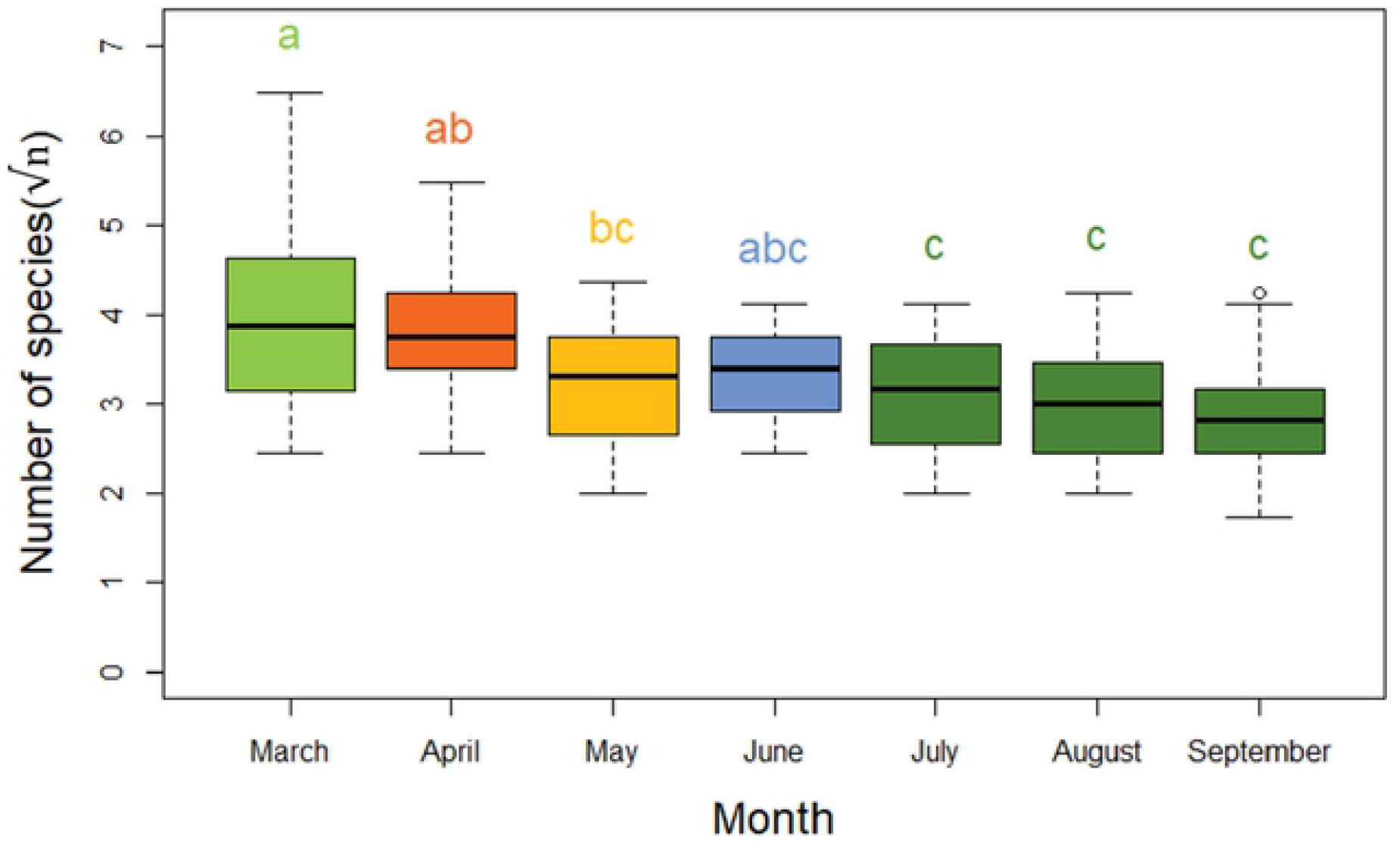
Boxplots of squared root taxa richness by sampling period. Letters and boxplot colours indicate significant differences according to Tukey’s test (*p* < 0.05).

NMDS displayed high variability in the composition of foraged pollen across the months and seasons (Fig 4). The greatest discontinuity separated spring (March, April, and May) from autumn (September). Concerning the floral composition foraged by the honeybees, May and August served as transition months to subsequent seasons. The permutation tests revealed that the month period (*F* = 6.87; *R*^2^ = 0.23; *p <* 0.001), site (*F* = 1.27; *R*^2^ = 0.1; *p <* 0.01), and landscape class (*F* = 2.01; *R*^2^ = 0.03; *p <* 0.001) were significant explanatory variables of pollen composition in the samples; however, the sampling period was attributed a larger proportion of the variance. From pairwise comparisons (i.e. letters from Fig 4), the urbanised sites hosted similar plant communities. Moreover, the structure of the plant communities varied significantly over the months until late summer and early autumn (i.e. August and September), when the floral composition harboured similar foraged plant communities.

**Fig 4.**
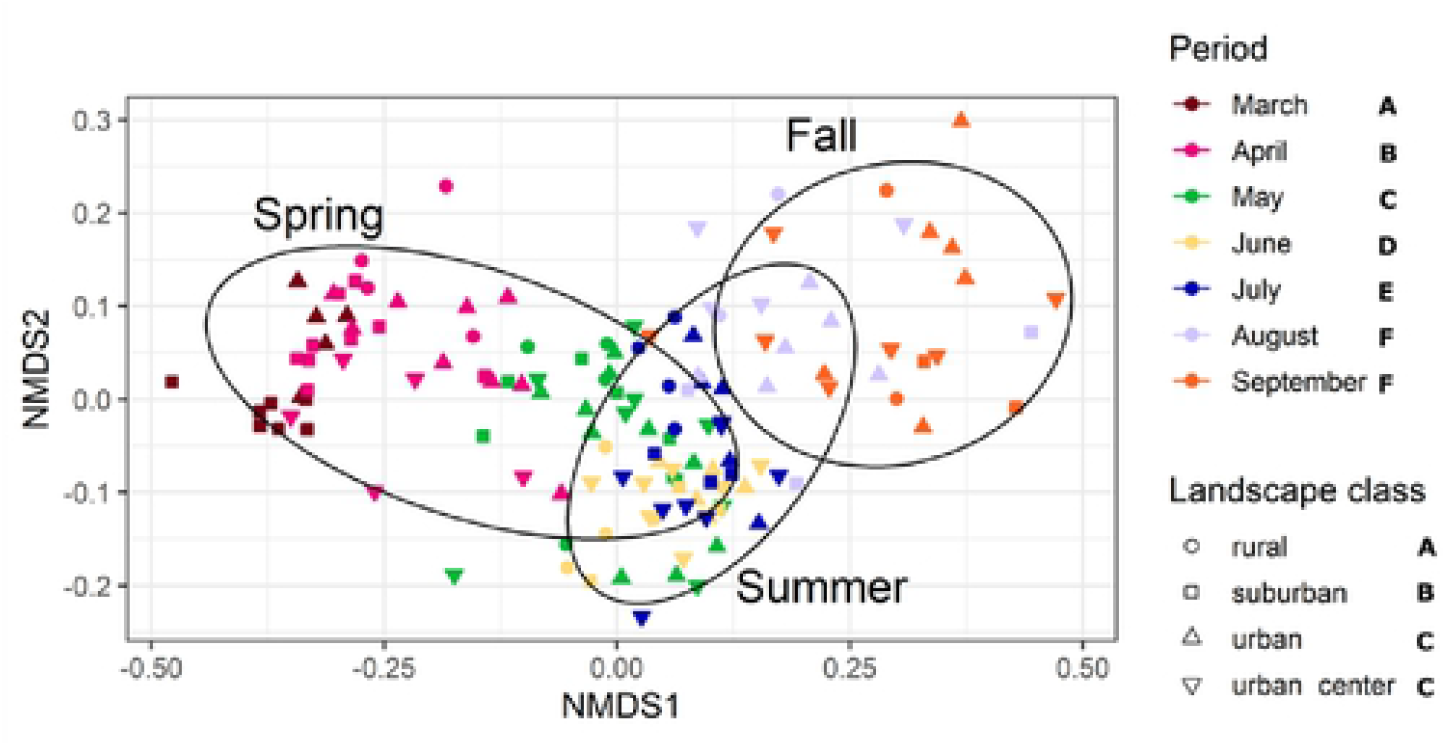
Non-metric multidimensional scaling (NMDS) of plant communities from the pollen incidence data. Dot shapes correspond to the landscape classes of pollen samples. Dot colours correspond to the sampling months, and the seasons are displayed by 80% prediction confidence ellipses. Letters indicate significant differences (*p* < 0.05) according to the pairwise post-hoc comparisons, with Bonferroni correction, of the foraged plant communities among the landscape gradient and the sampling period.

### Indicator species and trait-based analysis

The plant traits foraged by the honeybees varied significantly according to the nature (*G* = 99.0, *p <* 0.001) and native status (*G* = 69.1, *p <* 0.001) over the study months (Fig 5B), while only the plant strata showed significant differences (*G* = 10.7, *p <* 0.05) according to the landscape classes (Fig 5A).

**Fig 5.**
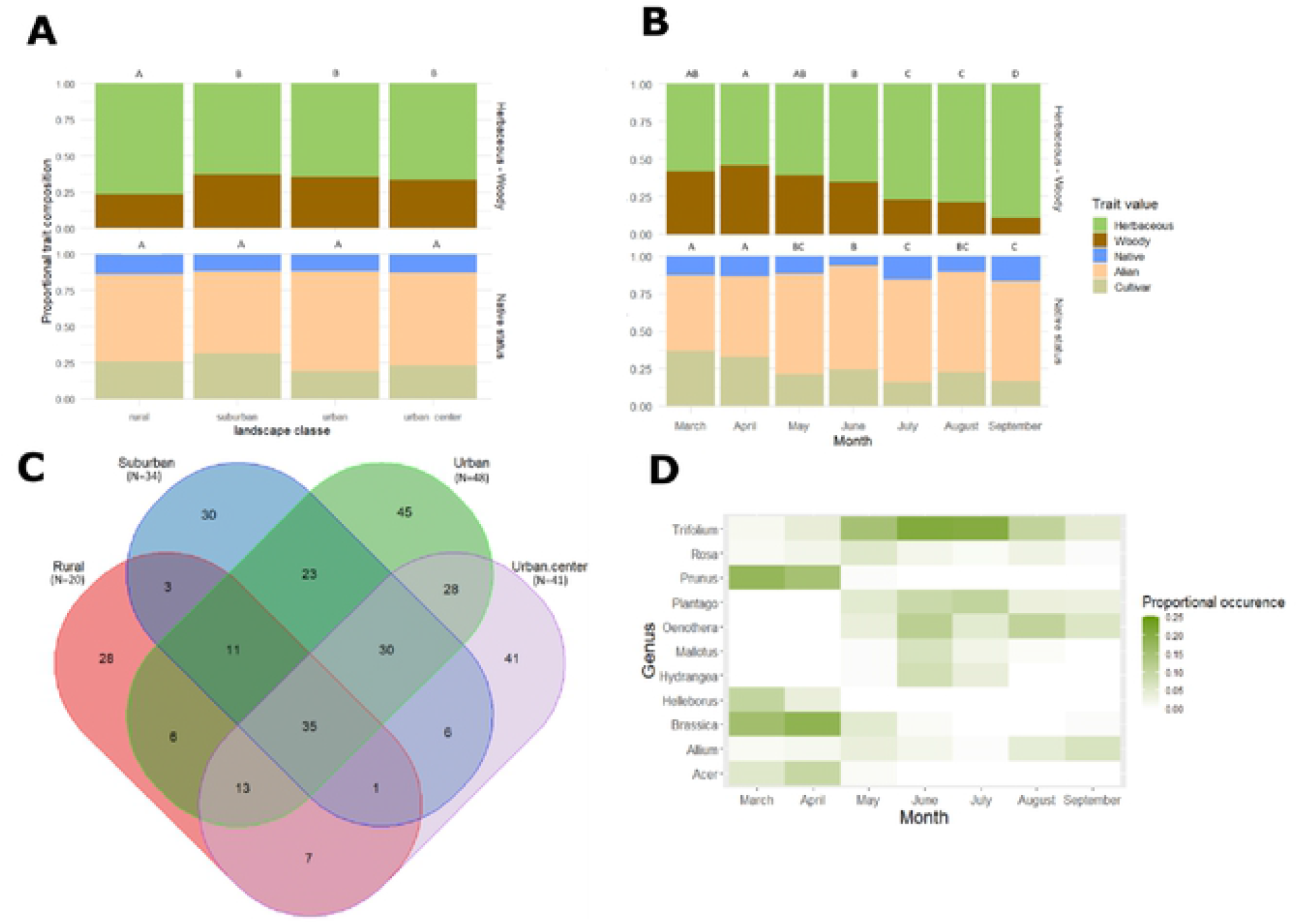
Proportional occurrences of the different plant traits, Venn diagram, and heatmap according to the landscape classes and the sampling months. (A) Proportional occurrences of the different plant traits (plant nature and nativity status) along the different landscape classes. Letters on top of the bar cluster homogeneous landscapes according to the significant results of post-hoc pairwise comparison with Bonferroni correction (*p* < 0.05). (B) Proportional occurrence of the different plant traits (plant nature and nativity status) across the sampling period. Letters on top of the bar cluster denote homogeneous sampling periods according to the significant results of post-hoc pairwise comparison with Bonferroni correction (*p* < 0.05). (C) Venn diagram indicating the overlap of plant taxa according to the different landscape classes. “*N*” corresponds to the number of samples per landscape. (D) Heatmap representing 11 most frequent genera (N = 62 taxa; 47% of the meta-barcoding dataset) ordered in descending order by their prevalence in all samples according to their monthly proportional occurrence. The proportional occurrence corresponds to the ratio of the number of observations for a genus per month to the number of samples for the specific month. The 11 genera were chosen according to the condition that their observation frequency is superior to 10% of the total occurrence of at least one season.

#### Effect of landscape

Significantly more herbaceous plant taxa structured the rural landscape (Fig 5A), even if plant composition traits were independent of landscape type. A total of 35 plant taxa were shared among all the landscapes over the sampling months, while 151 plant taxa were exclusively related to specific landscapes, corresponding to 27%, 22%, 24%, and 26% of the total plant taxa for rural, suburban, urban, and urban centre landscapes, respectively (Fig 5C). Plant families, such as Fabaceae, Rosaceae, Brassicaceae, Asteraceae, Plantaginaceae, and Onagraceae, were the most frequent taxonomic families encountered in all samples. However, their proportions varied according to the landscape (Fig S4). In urban and urban centre areas, leguminous plants prevailed in rural and suburban landscapes. However, the suburban landscape showed a higher frequency for the Brassicaceae, Ranunculaceae, and Rosaceae plant families. Surprisingly, anemophilous plants of the Poaceae family were more frequently foraged in the urbanised landscape than in the countryside (Fig S4).

#### Effect of sampling period

The proportion of foraged woody taxa decreased significantly over the seasons (*G* = 87.5, *p <* 0.001), with a peak of 46% in April and a low of 10% in September. Over the sampling months, honeybees foraged mainly on alien plant taxa (Fig 5B). Cultivar taxa were more foraged in spring than in the other two seasons (*G* = 32.9, *p <* 0.001). The most visited plant genera in March and April included *Prunus, Helleborus, Brassica*, and *Acer* taxa (Fig 5D). In contrast, four of the 11 most frequent genera emerged in late spring in May. Between April and June, a noticeable phenological turnover in the pollen composition (Fig 5D) was observed, with May serving as a transition bridge. This has already been highlighted by the discontinuities in the NMDS ordination (Fig 4). Following this shift, the genus *Trifolium* spp. was highly dominant in the June and July samples. In addition, the herbaceous genera *Plantago* and *Oenothera* spp. were also found in large proportions in combination with the woody genera *Mallotus and Hydrangea* spp. In August, the taxa from the genus *Oenothera* were the most represented with *Trifolium*, despite a reduction in its occurrence. A shift in pollen composition trends, with a reduction in highly proportional occurrence genera, was observed in August and September. In other words, plants detected in August and September were more distributed between the genera. Only the genus *Allium* showed an increase from August to September. Finally, *Trifolium* spp., *Rosa* spp., and *Allium* spp. were the only genera that were observed throughout the study period.

## Discussion

This study revealed interesting patterns of honeybee foraging habits along the urban-rural gradient throughout the 2019 active season among 17 different sites. In the present study, we used an unconventional approach, namely k-means clustering from landscape metrics, to categorise our sampling sites into four landscape classes. These landscape variables allowed the estimation of the effects of some ecological processes at the landscape level (i.e. foraging and plant dispersal) in assessing the diversity, connectivity, and aggregation of the patches [92,93]. Despite the convincing results of the grouping method, several reservations are worth mentioning. First, the selected foraging radius of 6 km accentuated the spatial autocorrelation issues on the landscape variables by increasing the foraging area overlaps among the sites [94]. This was not tested in the present study because we considered that there was no spatial influence on foraging preferences between each sampling colony due to the high variations in their foraging behaviour [95,96], even at the local scale for colonies of the same apiary [97]. Indeed, the foraging behaviour of honeybee colonies is mainly driven by: (i) the temporal colony needs as a function of its strength; (ii) the high density of pollen and nectar resources available near the colony; (iii) the capacity to make hourly colonial decisions for the most profitable flower patches; and (iv) the ability to tightly modulate its pollen reserves to buffer the colony from seasonal pollen breaks (e.g. long rainy periods, rarefaction of floral resources at the end of summer, etc.) [21]. Second, the 3-image resolution from satellites may result in some limitations, especially in complex landscapes, such as urban matrices. Therefore, this spatial resolution issue could be counterbalanced with the use of specific cameras, such as red-green-blue (RGB) or multispectral cameras, mounted on unmanned aerial vehicles to characterise floral identification and surfaces [98,99]. Nonetheless, this research domain is still in its infancy, particularly in the data processing of imagery classification by deep learning [100]. However, despite these spatial limitations, the approach led to satisfactory classification, which paves the way for further investigations.

The use of DNA meta-barcoding allowed for the identification of the pollen composition of the samples. This methods yields to the identification of 307 taxa, which is higher than previous studies owing to a greater sampling effort among sites and seasons [46,47,52]. This reflects a great diversity of plant resources and highlights foraging patterns, regardless of the landscape. Through our molecular analysis, we gauged potential biases that may arise from PCR amplification, sequencing errors, or arbitrary choices in the bioinformatics pipeline [101,102]. Moreover, recent studies aimed to compare pollen quantification from complex samples using a metagenomics approach against light microscopy. They recommended using a multi-locus meta-barcoding strategy in combining plastid loci, such as the *matK* or *trnL* markers with ITS2, to improve the correlation between the number of reads and classic palynology identification [103,104]. However, in the present study, we used one pair of ITS1 primers [74], which led to the consideration of all the generated datasets on their incidences due to the poor correlations between the number of sequence reads and visually identified pollen grains [36,41,105]. Furthermore, the robustness of the correlation depended primarily on the pollen counting methodology, DNA marker selection, plant taxonomy dataset, and number of PCR cycles [52,104]. Despite the need for further studies and improvements, this clearly provides a strong foundation for the study plant-pollinator interactions and the floral preferences of pollinating insects.

The landscape type did not influence forage plant richness, as reported in previous studies [49,51]. However, considering the spatial variation of taxa composition among the sites of our area of interest (i.e. beta diversity) [106], our results showed a spatial structure of foraged plant communities in countryside, suburban, and urban environments (i.e. by merging the urban and urban centre areas) (Fig 4). It is likely that honeybee colonies modify their foraging preferences due to the high prevalence of unattractive ornamental flowers in urban landscapes [14]. The urban matrix also offers smaller spread patches and less dense floral resources, which contribute to the foraging change of workers [47,97]. This shift in the prospected flora is also coupled to fulfil the nutritional demand with a diverse and complementary floral diet [107,108]. Therefore, it shows the importance of taking beta diversity and not only the local richness into account to understand the community structure of foraged plants throughout space and time scales to protect local biodiversity and support conservation planning [53,109]. Despite the landscape structure of the foraged plant community, the trait-based analysis revealed no significant preference for honeybee colonies, except for herbaceous plants in rural landscapes (Fig 5A).

In all the samples, 35 plant taxa were shared throughout all the landscapes studied, corresponding to 45% of all read counts. The top three plant families were Fabaceae, Rosaceae, and Brassicaceae in the samples (Fig S4). The presence of grasses (Poaceae) may seem surprising among the most frequent families in the samples (Fig S4), given that these plants are considered unsuitable for *A. mellifera* resource needs [110]. In view of their dominance, it seems unlikely that anemophilic pollen is the result of contamination by pollen blown from flowers onto the body of the bees, such as the rice paddy field in Japan [111]. Other recent studies have shown that pollinators (i.e. bees and syrphids), particularly *Apis* bees, interact with wind-pollinated plant species for their nutrient or nesting requirements [112]. This means that these grass species could be implemented in rural and urban management recommendations.

In contrast to the minor variations in landscape diversity, we observed a strong seasonal effect on plant richness (Fig 3), the foraged plant community, and traits (Fig 4 and 5B). We observed higher plant richness and foraged woody taxa in spring than in the other two seasons (Fig 3 and 5B), as reported previously [82,113,114]. The genera *Prunus* spp. and *Acer* spp. (Fig 5D) dominate the foraged woody stratum during this season, as these taxa constitute an important source of pollen for the early season [82,103,114,115]. Indeed, bee breads with high proportions of both genera were positively correlated with a higher protein content [107]. Particularly, the complexity and high range of foraged plants is known to be beneficial to the “nutritional value” of these bee bread stocks, and thus to honeybee immunity [28]. After the spring period, the proportion of foraged herbaceous strata gradually substituted woody taxa to reach approximately 90% of herbaceous foraged taxa in September, in agreement with previous studies [82,114]. This growing herbaceous stratum is mainly dominated by *Trifolium* spp. and *Plantago* spp. which, because of their long flowering period or floral consistency, explains the lower richness of taxa foraged in summer and autumn [114,116,117]. Moreover, clover species (i.e. *Trifolium* spp.) are highly ubiquitous in grasslands, such as meadows for rural areas or parks and gardens for urban areas [114,118] and may contribute to the concentration effect of amino acid content [107]. Concerning the temporal dynamics of the native traits (Fig 5B), the observations point to a decrease in the frequency of cultivars in favour of exotic taxa, which contradicts previous studies [119,120]. The highly anthropized and fragmented environments of the Tokyo region and its surroundings could explain the predominance of non-native floral species. However, this statement should be mitigated and requires further investigation, such as a complete plant inventory of the study site. Finally, we observed a transitional change from August by a rupture of the dominant flower prevalence, which may correspond to the seasonal dearth of floral resources (Fig 4 and 5B) [114,121]. From the perspective of the honeybees, they mitigate this effect by increasing their foraging range, requiring extra effort for sometimes worthless rewards [121]. Therefore, in an urban greening context, it would be relevant to ensure the diversity and abundance of floral resources near apiaries by targeting seasonal gaps in pollen collection [11]. Finally, the study of other co-variables, such as brood monitoring or estimating the pollen collection/reserve of each colony, could be used to compare the conditions of each sampled colony and improve our understanding of the foraging patterns of the colonies [122].

The availability of floral resources needs to be sufficient to host both domesticated honeybees and local wild pollinators. The percentage of impervious surfaces plays a major role in the pollination of biodiversity [123]. Therefore, populations of managed honeybees must be regulated to ensure wild pollinator populations are not adversely affected [18,124]. In contrast, green areas must be managed and well distributed to meet the demands of the pollinator community [125]. In summary, well-managed cities could play an active role in the preservation of insect pollinators, and thus provide hotspots for pollination services [126]. However, to do so, decision-makers will need to focus on regulating the introduction of honeybees (selection of native bee species, colony density, and control of pathogens and parasites) and on the availability of resources (proportion of impermeable surface area, melliferous plant species, and landscape diversity).

## Acknowledgments

The sequencing data have been deposited with links to BioProject accession number PRJNA74415 in the NCBI BioProject database (https://www.ncbi.nlm.nih.gov/bioproject/). This work was supported by JSPS KAKENHI (grant number 18KK0121) and the Yamada Research Grant. We thank Dr. Junko Kawai at Chiba University for his useful advice on DNA meta barcoding. The pollen was provided by following beekeepers in Japan: Ryuta Aikyo, Sayuri Aoki, Shinta Hegizono, Honami Kawabe, Tamaki Maruhashi, Haruko Nagae, Emi Nobunaga, Nobuyuki Okada, Takumi Sakou, Yuichi Shibasaki, Yosuke Toyomasu, Misa Uchino, and Tomoyuki Yuzurihara.

## Supplementary information

### Tables

**Table S1.** Climatic data of the study sites

| **Location** | **Prefectu** | **Elevation [m]** | **Mean Temperature [°C]** | **Precipitati** |
| --- | --- | --- | --- | --- |
| Enokisawa | Chiba | 14 | 15.6 | 1428 |
| Kuwata | Chiba | 23 | 15.6 | 1428 |
| Yachiyo | Chiba | 31 | 14.9 | 1394 |
| Ichihara | Chiba | 36 | 15.5 | 1550 |
| Nerima | Tokyo | 38 | 15.1 | 1448 |
| Shiba | Tokyo | 6 | 15.4 | 1442 |
| Togo | Tokyo | 29 | 15.4 | 1442 |
| Colombin | Tokyo | 24 | 15.4 | 1442 |
| Shinjyuku | Tokyo | 32 | 15.4 | 1442 |
| Toyosu | Tokyo | 6 | 15.4 | 1442 |
| Yamatecho | Kanagaw | 23 | 15.6 | 1554 |
| Ishikawach | Kanagaw | 29 | 14.7 | 1488 |
| Gumyoji | Kanagaw | 13 | 15.6 | 1554 |
| Honmoku | Kanagaw | 12 | 15.6 | 1554 |
| Nishichiba | Chiba | 17 | 15.3 | 1435 |
| Kashiwano | Chiba | 19 | 14.7 | 1358 |
| Inohana | Chiba | 17 | 15.3 | 1435 |

### Supplementary Figure legends

**Fig S1.**
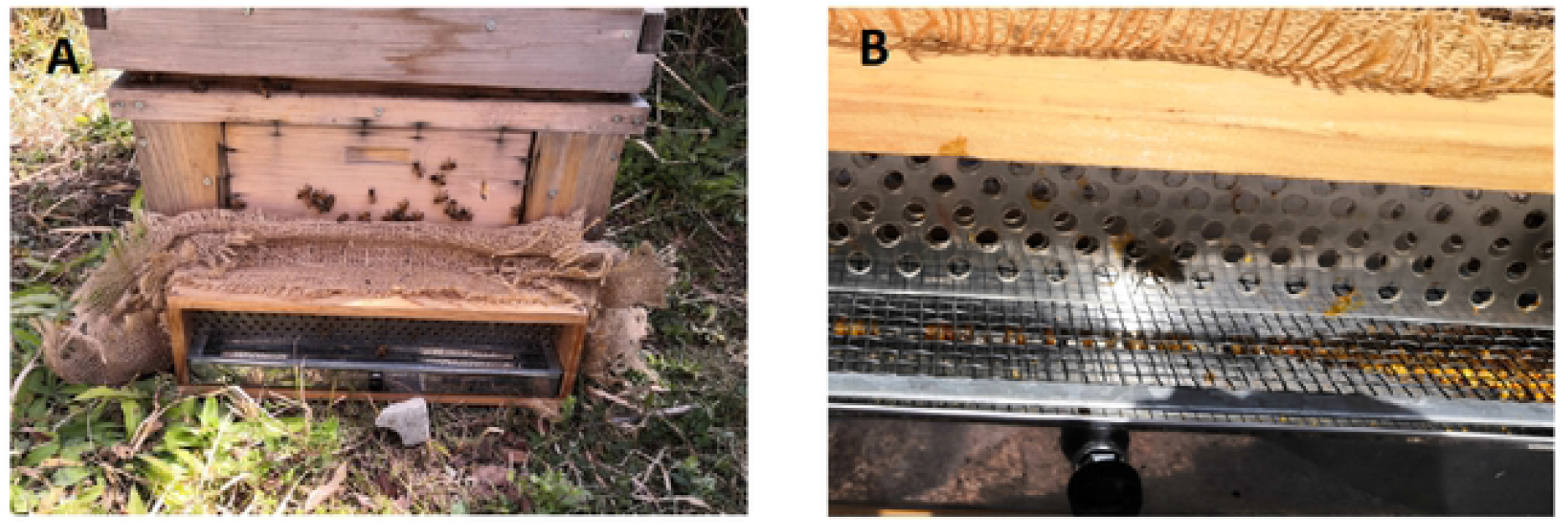
Pollen traps. (A) Pollen trap set-up at the entrance of the selected hive in the Nishi-Chiba campus of the Chiba University (25 March 2020). (B) Close-up of pollen trap with pollen balls collected in the trail in the Nishi-Chiba campus of the Chiba University (17 June 2020).

**Fig S2.**
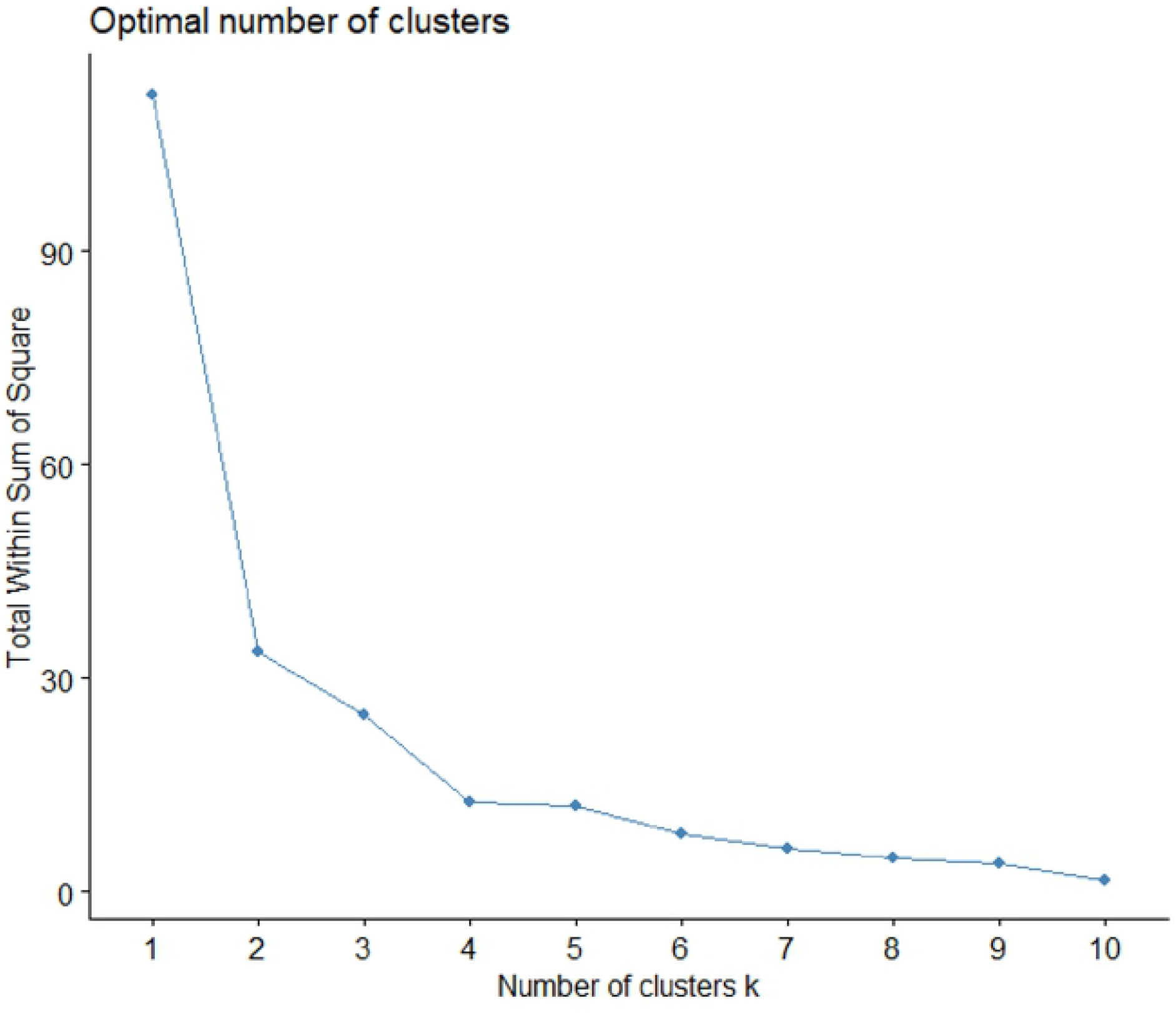
Elbow clustering plot. Elbow method plotting the total within sum of square explained in function of the number of k clusters. The elbow of the curve suggests the number of groups to retain for k-means clustering analysis.

**Fig S3.**
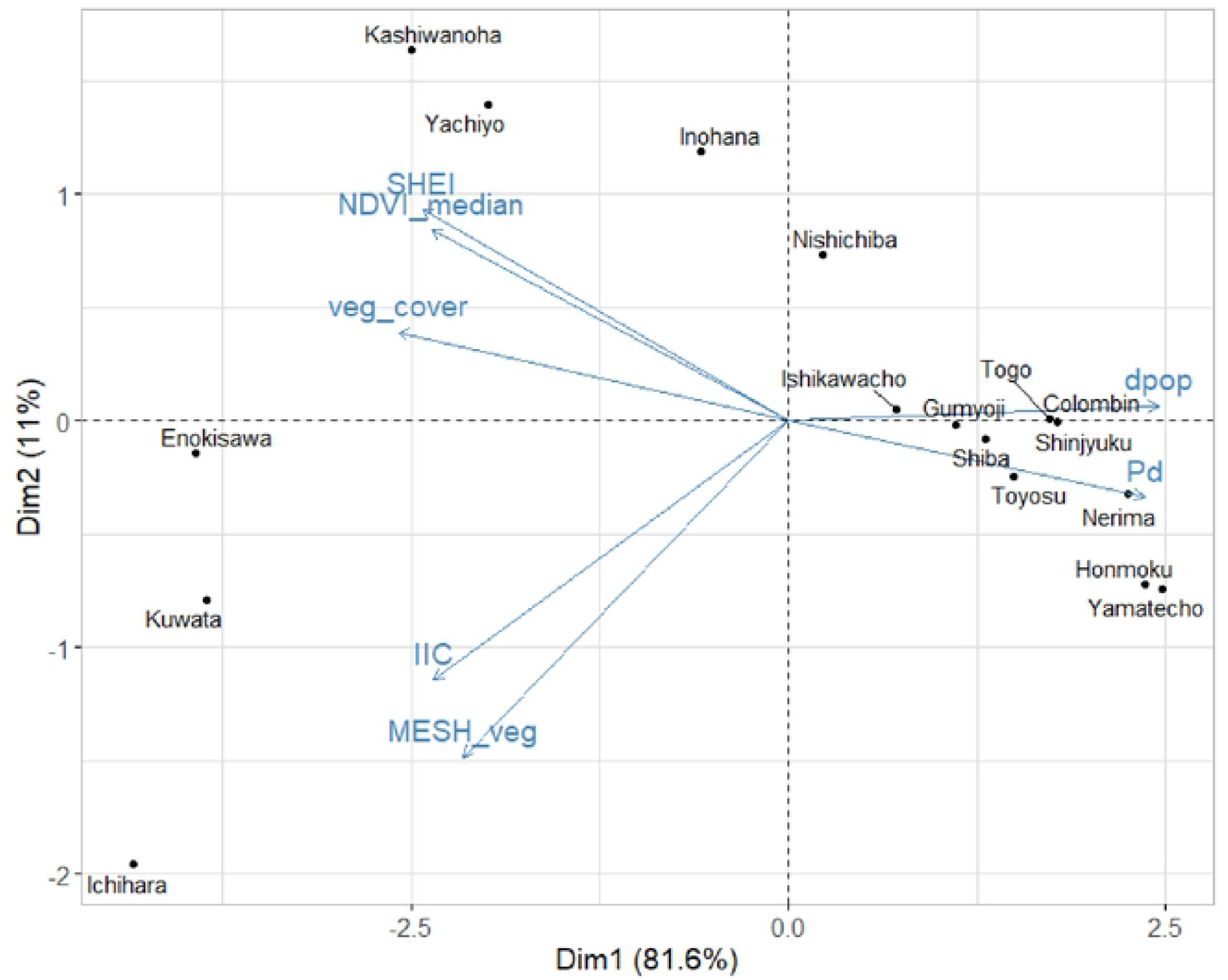
Principal component analysis (PCA) biplot of individual sites and landscape variables. The landscape variables (in blue) are Pd: Patch density [patches/km^2^], dpop: demographic density (number of inhabitant/km^2^), MESH_veg: effective mesh size of vegetation [-], IIC: Integral index of connectivity [-], veg_cover: vegetation cover (%), NDVI_median: median of the NDVI of the cells superior to 0.2 [-], and SHEI: Shannon’s evenness index [-].

**Fig S4.**
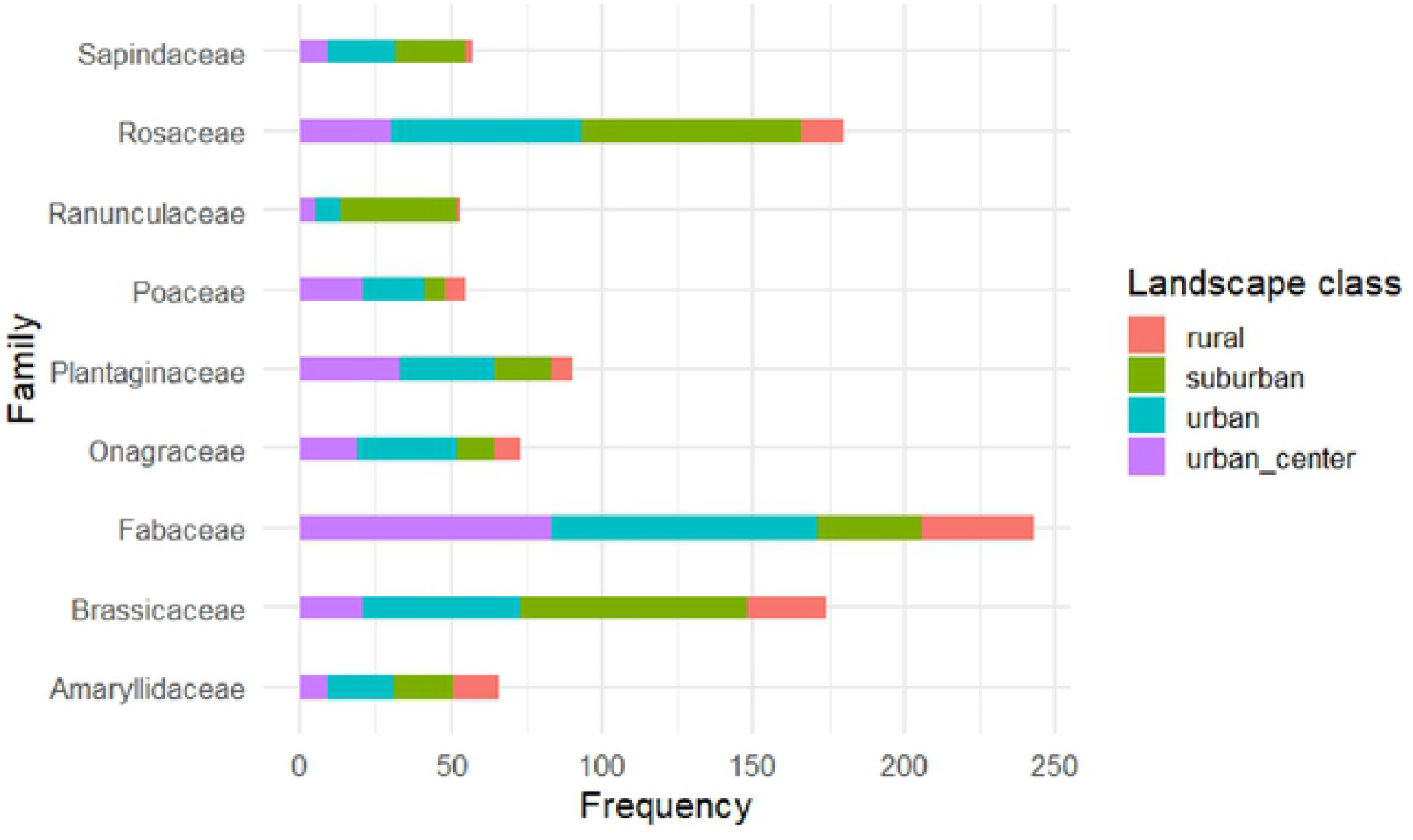
Bar plot showing the nine most frequent families observed in the samples in contrast with the landscape classes. Colour proportions correspond to the landscape classes.

